# Toxoplasmosis, but not borreliosis, is associated with psychiatric disorders: A cross-sectional survey on 46 thousand of subjects

**DOI:** 10.1101/231803

**Authors:** Jaroslav Flegr, Jiří Horáček

## Abstract

Infection by the parasite *Toxoplasma*, which affects about 33% of world population, is associated with increased risk of several mental health disorders, the most strongly with schizophrenia. It is unknown whether toxoplasmosis really plays a substantial role in the etiopathogenesis of these disorders and whether schizophrenia is associated with this infection the most strongly, or whether this association has just been the most intensively studied for historical reasons. We used the data from 3,440 and 7,800 subjects tested for toxoplasmosis and borreliosis, respectively, who took part in an internet survey, for searching in the associations of these infections with 22 mental health disorders and other indices of impaired mental health. The typical symptom associated with toxoplasmosis was anxiety, and the typical toxoplasmosis-associated disorders were autism (OR=4.86), schizophrenia (OR=3.34), attention deficit hyperactivity disorder (OR=3.02), Asperger syndrome (OR=2.49), antisocial personality disorder (OR=1.81), OCD (OR=1.69), and anxiety disorder (OR=1.57). Borreliosis was associated only with symptoms of depression and with major depression (OR=1.65). The negative effects of borreliosis were detectable only in the *Toxoplasma*-infected subjects. Toxoplasmosis could play a substantial role in the etiopathogenesis of mental health disorders and its association with schizophrenia is the second strongest association, after autism.

## 1 Introduction

About one third of the world population is infected with the coccidian parasite *Toxoplasma gondii*. The course of postnatally acquired toxoplasmosis in immunocompetent subjects is mild and therefore the so-called latent toxoplasmosis has been mostly considered as clinically insignificant. However, the results of recent studies show that this picture could be wrong. *Toxoplasma*-seropositivity is associated with the increased risk of many disorders (Flegr and Escudero, 2016) and between-country differences in seroprevalence of toxoplasmosis could explain 23% of the total variability in disease burden in European countries (Flegr et al., 2014). The connection between certain mental health disorders, especially schizophrenia and toxoplasmosis, has been confirmed to exist behind any reasonable doubt, for review see (Sutterland et al., 2015; Torrey et al., 2012). Moreover, the causal role of toxoplasmosis in the development of schizophrenia has been confirmed by a longitudinal study (Niebuhr et al., 2007). It has been even documented that the changes in brain morphology that are characteristic for schizophrenia, such as grey matter reduction in frontal and temporal cortices, caudate, median cingulate, and thalamus, are in fact typical for *Toxoplasma*-seropositive schizophrenia patients (Horacek et al., 2012). Congruently, the *Toxoplasma*-seropositive patients express more prominent positive symptoms of schizophrenia (Holub et al., 2013; Wang et al., 2006) and have a 15-times higher probability of having a continuous course of disease than the *Toxoplasma*-free patients (Celik et al., 2015). Far fewer studies have shown the association of toxoplasmosis with other mental health disorders. About ten studies showed the association of toxoplasmosis with bipolar disorder, and less than five with obsessive compulsive disorder, learning disorder, autism, and anxiety disorder; for reviews see (Flegr, 2015a; Sutterland et al., 2015).

*Toxoplasma* is just one of many neurotropic pathogens that infect a large part of the human population in many developed countries. It can therefore be speculated as to whether this particular parasite really plays the prominent role in the etiology of mental health disorders, or whether this parasite is just studied more often, for example in the context of its well established manipulation activity aimed to increase the chance of transmission from the intermediate to definitive host (the cat) by predation (Berdoy et al., 2000; Moore and Gotelli, 1996). The intensive study of the toxoplasmosis-schizophrenia association could be just a historical accident as the parasite hypothesis of schizophrenia has been systematically and intensively studied by the teams of Fuller Torrey and Robert Yolken at Stanley Research Institute since early 2000s (Yolken et al., 2001).

To test whether *Toxoplasma gondii* plays a prominent role in the etiology of mental health disorders, we compared the effect of *Toxoplasma*-seropositivity with the effect of *Borrelia*-seropositivity. It was documented (Hajek et al., 2002) that *Borrelia*-seropositivity is more prevalent in psychiatric patients (33%) than among matched healthy subjects (19%). Both pathogens have a similar seroprevalence in the Czech population and many inhabitants of Czechia are being tested for the presence of the anamnestic antibodies against either the protozoan *Toxoplasma* or the bacterium (spirochete) *Borrelia*.

It is not known, whether the association of *Toxoplasma* with schizophrenia is the strongest or whether it is just most often studied. To address this question, we performed the systematic search for any association between *Toxoplasma* or *Borrelia*-seropositivity and the 22 most prevalent mental health disorders. However, the incidence of certain disorders, e.g. schizophrenia, is relatively low and doing many statistical tests requires performing the rigorous correction for multiple tests. For these reasons, we needed to analyze a very large data set. To this end, we utilized data from a recently obtained, large internet-based cohort designed for evolutionary psychological, psychopathological and parasitological studies that was primarily aimed at searching for possible effects of biological and nonbiological factors on human sexual preference and behavior. This study was very popular in Czechia and the corresponding electronic questionnaire was completed by nearly 50 thousand of subjects (about 0.5 % of all inhabitants of Czechia), many of them tested for either toxoplasmosis or borreliosis.

## 2 Materials and methods

### 2.1 Study population

The data was originally collected for the purpose of another study (Flegr and Kuba, 2016). The subjects were invited to participate in the study using a Facebook-based snowball method (Kankova et al., 2015) by advertisements published in various papers and electronic media, as well as TV and radio broadcasting. The invitation to participate in a “study testing certain evolutionary psychological and parasitological hypotheses, containing many questions related to sexual life” was also posted on the wall of the Facebook page “Guinea pigs” (“Pokusní králíci” in Czech) for Czech and Slovak nationals willing to take part in diverse ethological and psychological projects (www.facebook.com/pokusnikralici). The participants were informed about the aims of the study on the first page of the electronic questionnaire. They were also provided with the following information: “The questionnaire is anonymous and the obtained data will be used exclusively for scientific purposes. Your cooperation in the project is voluntary and you can terminate it at any time by closing this web page. You can also skip any uncomfortable questions; however, the most valuable data is complete data. Only subjects above fifteen years old are allowed to take the questionnaire. If you agree to participate in the research and are above 15, press the “Next” button. Some pages of the questionnaire contained the Facebook share button. These buttons were pressed by 1,319 participants, which resulted in obtaining data from about 46,000 respondents in total between 22^nd^ January 2015 and 8^th^ March 2017. The project, including the method of obtaining an electronic consent to participate in the study, was approved by the Ethical committee of the Faculty of Science, Charles University (No. 2015/01).

### 2.2 Questionnaires

The primary aim of present project was to test certain hypotheses from the fields of evolutionary psychology, psychopathology, sexology and parasitology. Therefore, the electronic survey consisted of 5 already published questionnaires studying various facets of human sexuality, such as The Hurlbert Index of Sexual Narcissism (Hurlbert, 1994), the Sexual Attitude Scale SOI-R (Penke and 2008), the Three-Domain Disgust Scale (Tybur, 2009), the International Personality Item Pool - Dominance scale (IPIP) (Goldberg, 1999; Goldberg et al., 2006), and the Attraction to sexual aggression scale, (Malamuth and 1989) modified and supplemented with questions to cover a broader spectrum of sexual preferences and sexual behavior. The survey also contained an anamnestic questionnaire collecting various socioeconomic, demographic, health related, epidemiologic, and psychological data and three projective psychological tests. Altogether, the survey consisted of more than 700 questions and the mean time necessary to complete it was about 110 minutes (the mode was 97 minutes). In the present study, we used only the information about gender, age, mental health-related variables and *Toxoplasma* and *Borrelia* infections. The subjects were also asked to rate intensity of any psychiatric problems diagnosed by doctors and undiagnosed by doctors using two graphic 0-100 scales, and then to check which mental health disorders they suffer (both diagnosed and undiagnosed by a clinician) on the list of 24 disorders, see the Table 2. Only two subjects tested for *Toxoplasma* and three tested for *Borrelia* reported to suffer from Parkinson‘s disease and only two tested for *Toxoplasma* and six tested for *Borrelia* reported to suffer from Alzheimer’s disease; therefore the associations between the infections and these two disorders were not analyzed. Then, they were asked to rate the intensity of suffering from particular neuropsychiatric symptoms (depression, mania, phobias, anxiety, and obsessive behavior) by moving sliders on graphic scales 0-100, anchored with Newer (0) and Intensively or frequently (100). The subjects have been also asked whether they are *Toxoplasma*- or *Borrelia*-infected. They were reminded that *Toxoplasma* is ‘‘a parasite of cats, dangerous especially to pregnant women’’ and *Borrelia* is “the microbe transmitted by ticks for which no vaccine is available”. (People in Czechia are often vaccinated against viral tick-borne encephalitis, however, no commercial vaccine against European strains of *Borrelia* is now available.) The response ‘‘I do not know, I am not sure’’ was set as a default answer which the respondents could change by selecting either ‘‘No, I was tested by a doctor and the result of my laboratory tests was negative’’ or ‘‘Yes, I was tested by a doctor and I had antibodies against *Toxoplasma* (against *Borrelia*)’. The respondents to our questionnaires had three options: they could complete any questionnaire absolutely anonymously, they could sign the finished questionnaire by a code obtained after the anonymous registration, or they could sign the finished questionnaire by a code obtained after the non-anonymous registration.

### 2.3 Statistical methods

Before statistical analysis, less than 1% of suspicious data (too high or too short body height, too low or too high body mass or age, too short duration of the test, etc.) were filtered out. Two subjects were also excluded because they checked nearly all mental health disorders, including Parkinson‘s and Alzheimer‘s disease.

Statistica v.10.0. was used for all statistical tests. Differences in the prevalence of particular disorder between men and women have been tested with logistic regression with sex, age and *Toxoplasma* and *Borrelia* seropositivity as the predictors. The effects of infections on continuous or ordinal variables (like intensity of suffering from anxiety or number of different mental health disorders were measured with ANCOVA. However, nonparametric tests (partial Kendall correlation test with age as covariate) provided qualitatively equivalent results (not shown). False discovery rate (preset to 0.20) was controlled with the Benjamini-Hochberg procedure (Benjamini and Hochberg, 1995). The raw data set is available at: 10.6084/m9.figshare.5593288.

## 3 Results

The final data set contained the information by 18,428 women (mean age 31.6, SD =12.1) and 27,458 men (mean age 35.9, SD =13.1); the difference in age between both sexes was significant (t_4584_=-35.9, p<0.0005). The information about mental health status provided by 10,956 women (mean age 30.5, SD =10.9) and 12,268 men (mean age 34.8, SD =12.0); the difference in age between men and women was highly significant (t_23222_=-28.5, p<0.0005). Among these subjects, 2,354 women (19.3% seropositive) and 1,361 men (11.1% seropositive) provided the information about their toxoplasmosis status and 3,252 women (23.0% seropositive) and 3 865 men (19.8% seropositive) provided the information about their borreliosis status. Only 71 (2.2%) women and 32 (0.83%) men reported both *Toxoplasma*- and *Borrelia*-seropositivity. The subjects who reported to be *Toxoplasma*-seropositive had a relatively higher probability of reporting also *Borrelia*-seropositivity (women: 33.8% vs 18.7%, Chi^2^=24.20, p<0.0005; men: 35.0% vs 9.6%, Chi^2^=84.4, p<0.0005). Using logistic regression with toxoplasmosis, female sex, and age as predictors, the data showed that all of these factors had significant effects on the probability of being *Borrelia*-seropositive (toxoplasmosis OR=3.10, p<0.0005; female sex OR=0.56, p<0.0005; age OR_range_=3.15, p<0.0005).

ANCOVA tests with independent variables of age, sex and either toxoplasmosis or borreliosis showed that Toxoplasma-seropositive subjects, especially women, reported more serious problems with depression (p=0.042), and anxiety (p=0.003), while no effect of *Borrelia*-seropositivity was significant. The results of the larger model that included age, sex and both toxoplasmosis and borreliosis as the predictors computed for the subset of 1997 raters reporting the results of the serological test for both toxoplasmosis and borreliosis were more informative, see the table 1 and Fig. 1.

**Table 1.** Effects of *Toxoplasma-* and *Borrelia-seropositivity* on reported psychiatric symptoms

|  | depression |  | mania |  | phobia |  | anxiety |  | obsessions |  |
| --- | --- | --- | --- | --- | --- | --- | --- | --- | --- | --- |
|  | p | eta<sup>2</sup> | p | eta<sup>2</sup> | p | eta<sup>2</sup> | p | eta<sup>2</sup> | p | eta<sup>2</sup> |
| age | **0.000** | 0.008 | **0.000** | 0.030 | **0.000** | 0.018 | **0.000** | 0.017 | **0.000** | 0.030 |
| sex | **0.000** | 0.013 | 0.340 | 0.001 | **0.000** | 0.018 | **0.000** | 0.023 | 0.213 | 0.001 |
| toxoplasmosis | **0.004** | 0.004 | 0.922 | 0.000 | 0.285 | 0.001 | **0.000** | 0.007 | **0.026** | 0.003 |
| borreliosis | 0.054 | 0.002 | 0.416 | 0.000 | 0.380 | 0.000 | 0.039 | 0.002 | 0.830 | 0.000 |
| toxopl. x borrel. | 0.136 | 0.001 | 0.903 | 0.000 | 0.355 | 0.001 | 0.080 | 0.002 | 0.865 | 0.000 |

The $p$ value $< 0.00005$ is coded as 0.000; the associations significant after the correction for multiple (five) tests are printed in bold.

**Fig. 1.**
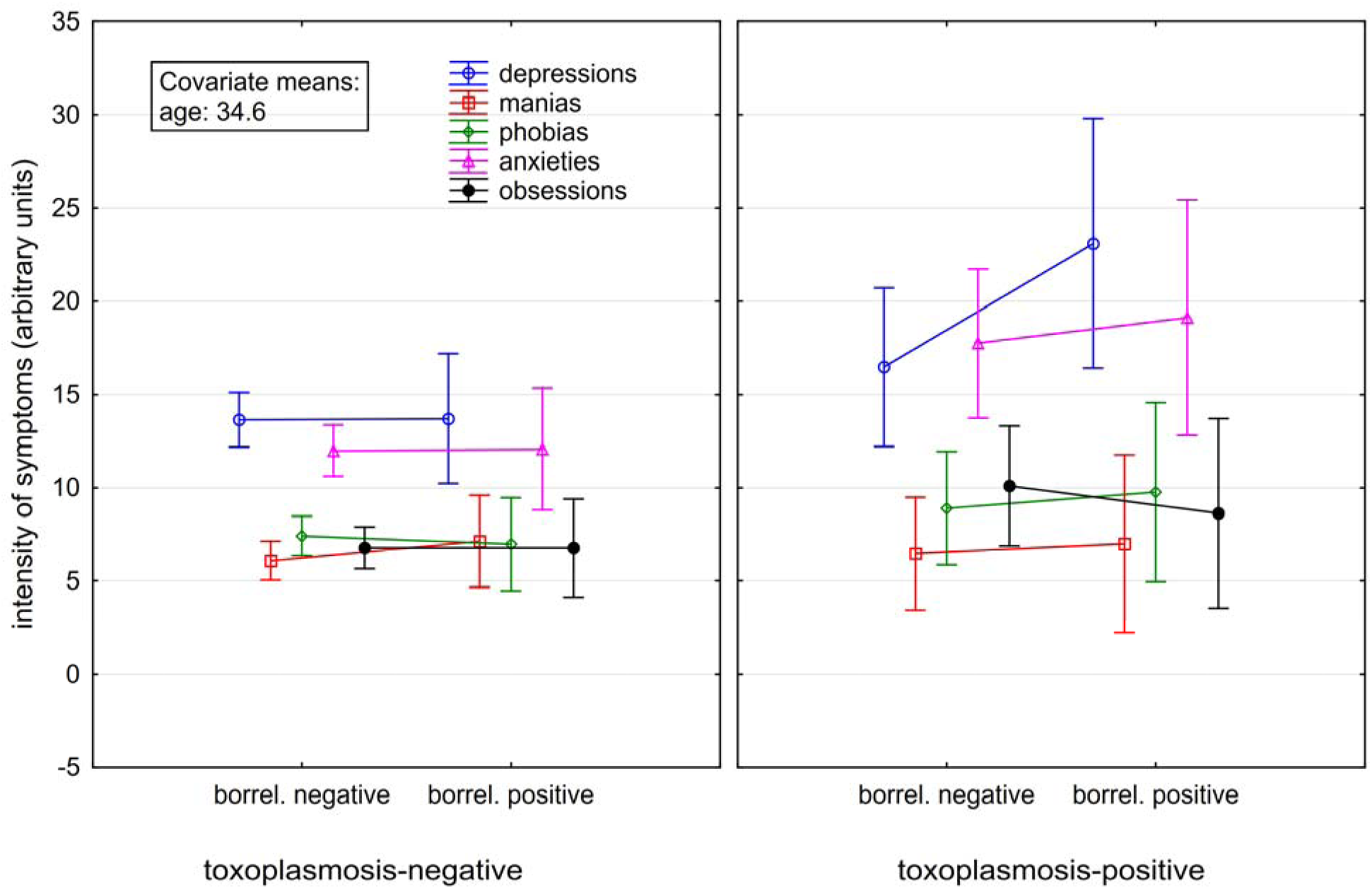
Effects of *Toxoplasma*- and *Borrelia*-seropositivity on reported psychiatric symptoms. *The spreads show 95% confidence intervals.*

After the correction for multiple tests, toxoplasmosis had a significant effect on the intensity of symptoms of depression, anxiety, and obsessions. Borreliosis had a significant effect on anxiety, (p 0.037) but this finding did not survive correction for multiple tests, and a borderline effect on depression (p=0.054). The Fig. 1 shows that borreliosis was associated with depressive and anxiety symptoms only in *Toxoplasma*-seropositive subjects; formally, however, the effects of the toxoplasmosis-borreliosis interaction on anxiety (p=0.080) and depression (p=0.136) were not significant.

ANCOVA analyses with sex, age, and toxoplasmosis as predictors showed that toxoplasmosis positively correlated with the intensity of suffering from diagnosed mental health disorders (p=0.001, eta^2^=0.004), the intensity of suffering from mental health disorders undiagnosed by medical doctors (p=0.003, eta^2^=0.003), and with the number of different mental health disorders respondents checked on the list (p<0.0005, eta^2^=0.005). These associations were weaker or absent for borreliosis: the intensity of suffering from diagnosed mental health disorders (p=0.005, eta^2^=0.001), the intensity of suffering from undiagnosed mental health disorders (p=0.796, eta^2^<0.0005), and the number of different mental health problems (p=0.121, eta^2^<0.0005). In the *Toxoplasma*-free subjects, the number of mental health disorders in *Borrelia*-free and *Borrelia*-infected subjects did not differ while in the *Toxoplasma-infected* subjects this number was about two times higher in *Borrelia*-infected subjects (toxoplasmosis-borreliosis interaction: p=0.013), see the Fig. 2. The tables 2 and 3 show the association of *Toxoplasma* and *Borrelia* infections with the incidence of particular mental health disorders.

**Fig. 2.**
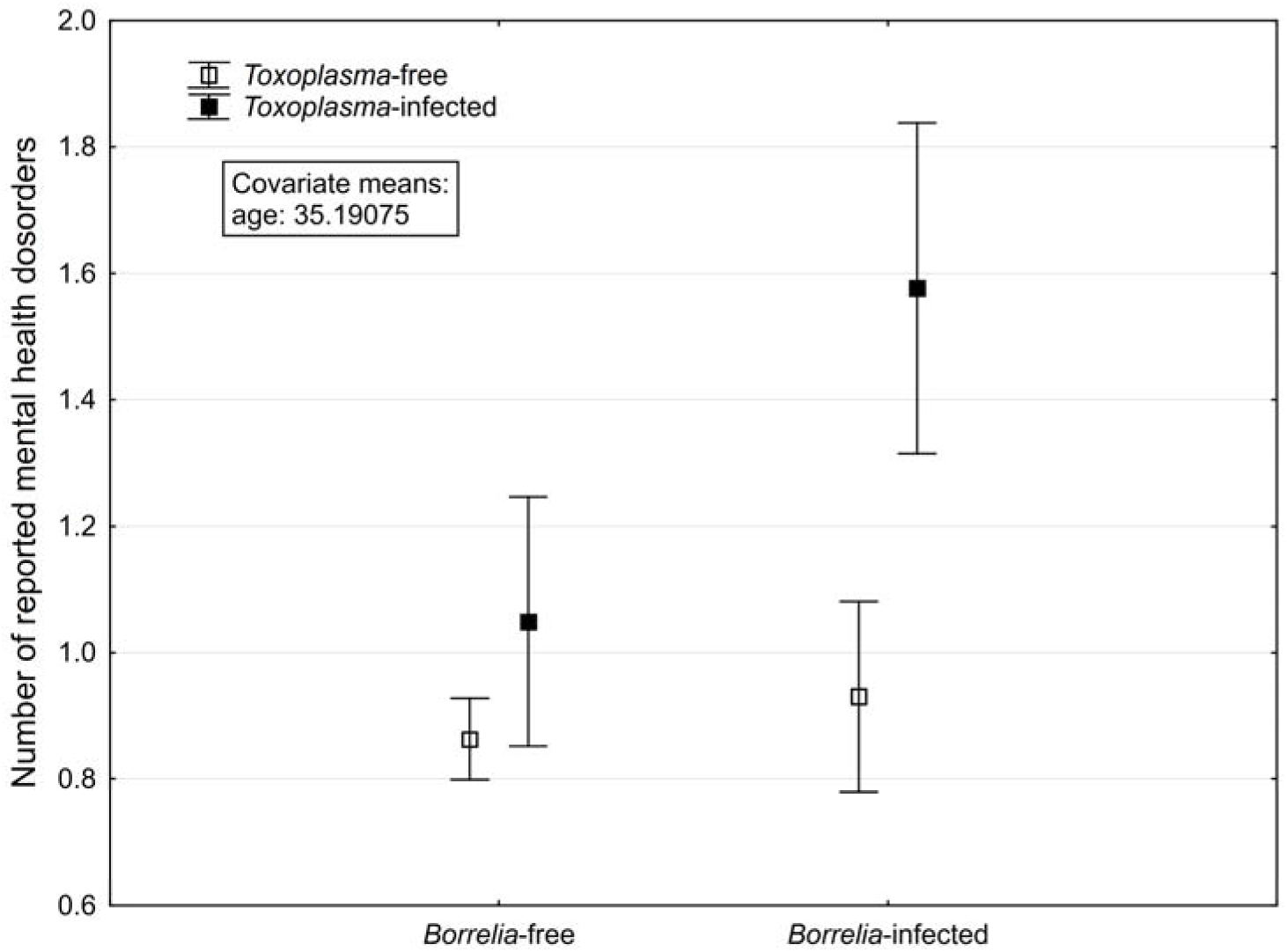
Association of the number of reported diagnosed and undiagnosed mental health disorders with *Toxoplasma* and *Borrelia* infections. *The spreads show 95% confidence intervals.*

**Table 2.**
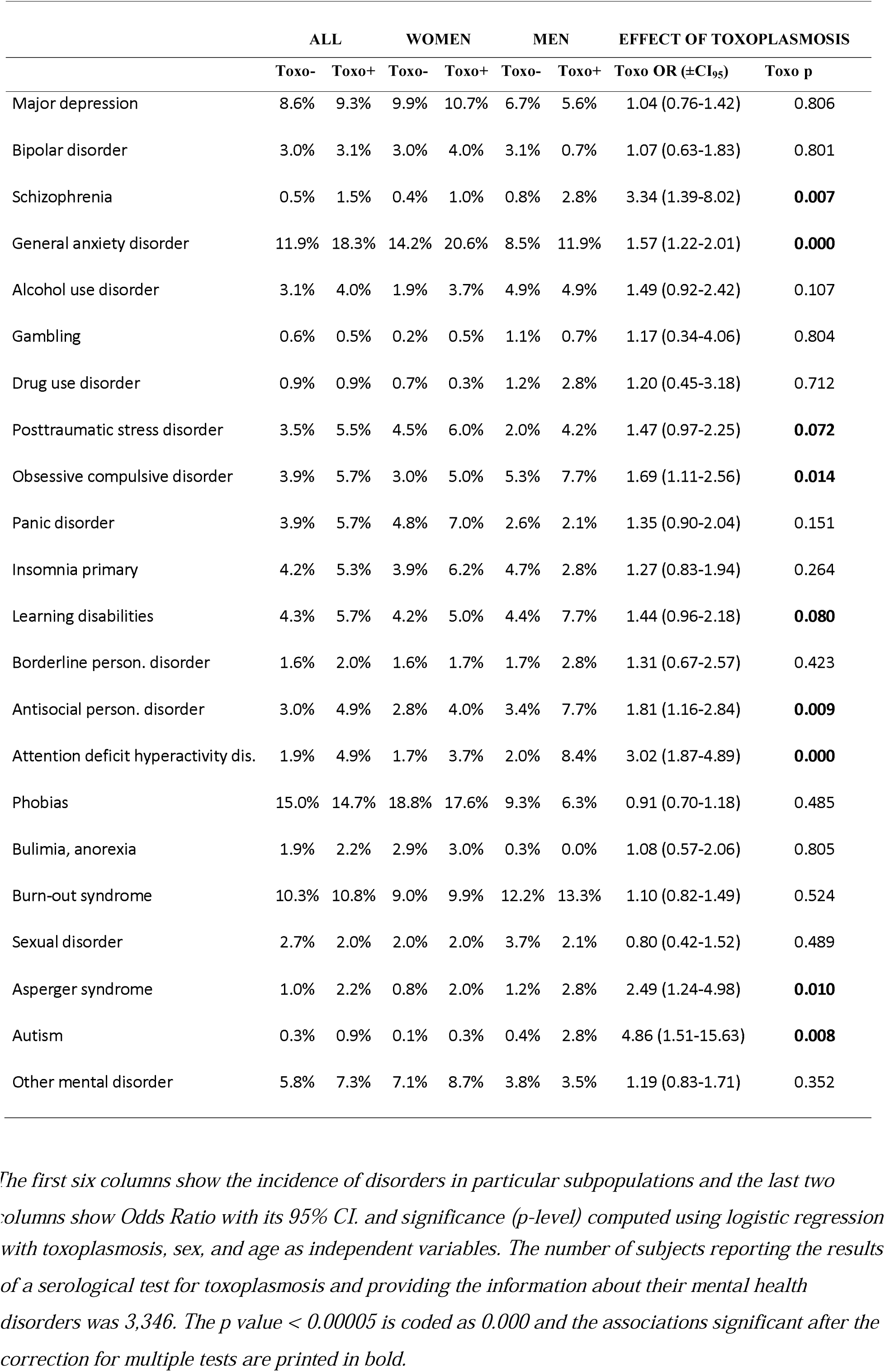
Incidence of particular mental disorders in *Toxoplasma*-seropositive and *Toxoplasma*-seronegative subjects and their association with toxoplasmosis measured with logistic regression

**Table 3.** Incidence of particular mental disorders in *Borrelia*-seropositive and *Borrelia*-seronegative subjects and their relation measured with logistic regression

|  | ALL |  | WOMEN |  | MEN |  | EFFECT OF BORRELIOSIS |  |
| --- | --- | --- | --- | --- | --- | --- | --- | --- |
| | Borrel- | Borrel+ | Borrel- | Borrel+ | Borrel- | Borrel+ | Borrel OR (±CI<sub>95</sub>) | Borrel p |
| Major depression | 7.8% | 10.3% | 9.8% | 12.9% | 6.1% | 7.6% | 1.65 (1.14-2.38) | **0.008** |
| Bipolar disorder | 2.8% | 2.4% | 3.2% | 2.6% | 2.5% | 2.3% | 1.15 (0.63-2.12) | 0.645 |
| Schizophrenia | 0.7% | 0.7% | 0.9% | 0.9% | 0.6% | 0.5% | 1.86 (0.53-6.50) | 0.329 |
| Anxiety disorder | 13.8% | 13.9% | 18.5% | 17.3% | 9.7% | 10.5% | 1.24 (0.90-1.70) | 0.193 |
| Alcohol use disorder | 3.8% | 4.2% | 2.4% | 2.6% | 5.0% | 6.0% | 1.14 (0.65-2.01) | 0.640 |
| Gambling | 0.6% | 0.8% | 0.3% | 0.2% | 0.9% | 1.4% | 0.41 (0.05-3.36) | 0.409 |
| Drug use disorder | 1.6% | 1.3% | 1.4% | 0.7% | 1.7% | 1.9% | 1.58 (0.61-4.11) | 0.349 |
| Posttraumatic stress disorder | 3.2% | 4.6% | 5.0% | 6.7% | 1.7% | 2.4% | 1.64 (1.01-2.65) | 0.045 |
| Obsessive compulsive disorder | 4.8% | 4.9% | 4.5% | 5.0% | 5.0% | 4.9% | 1.08 (0.62-1.86) | 0.794 |
| Panic disorder | 4.5% | 4.6% | 6.5% | 6.1% | 2.8% | 3.1% | 0.76 (0.43-1.36) | 0.359 |
| Insomnia primary | 4.7% | 4.5% | 5.5% | 4.5% | 4.1% | 4.4% | 1.03 (0.63-1.68) | 0.919 |
| Learning disabilities | 5.6% | 5.5% | 5.1% | 4.8% | 6.0% | 6.2% | 1.32 (0.81-2.16) | 0.270 |
| Borderline person. Disorder | 2.0% | 1.2% | 2.5% | 1.3% | 1.6% | 1.1% | 1.12 (0.50-2.50) | 0.781 |
| Antisocial pers. Disorder | 4.6% | 4.3% | 4.5% | 2.7% | 4.8% | 6.1% | 1.22 (0.68-2.19) | 0.503 |
| Attention deficit hyperactivity dis. | 3.3% | 3.5% | 3.1% | 3.5% | 3.5% | 3.6% | 1.22 (0.63-2.36) | 0.555 |
| Phobia | 15.9% | 15.6% | 22.0% | 21.1% | 10.6% | 10.0% | 1.22 (0.89-1.66) | 0.218 |
| Bulimia, anorexia | 2.3% | 1.9% | 4.5% | 3.6% | 0.4% | 0.2% | 0.70 (0.30-1.60) | 0.392 |
| Burn-out syndrome | 11.4% | 11.9% | 10.2% | 11.9% | 12.4% | 11.9% | 1.25 (0.89-1.76) | 0.190 |
| Sexual disorder | 3.3% | 3.2% | 2.5% | 2.1% | 4.0% | 4.4% | 1.18 (0.61-2.28) | 0.621 |
| Asperger syndrome | 1.2% | 1.5% | 0.9% | 1.3% | 1.5% | 1.8% | 1.27 (0.49-3.30) | 0.617 |
| Autism | 0.3% | 0.3% | 0.2% | 0.2% | 0.4% | 0.4% | 1.60 (0.28-8.96) | 0.595 |
| Other mental disorder | 4.7% | 6.2% | 6.1% | 8.9% | 3.5% | 3.3% | 1.66 (1.08-2.56) | 0.021 |

The first six columns show the incidence of disorders in particular subpopulations and the last two columns show Odds Ratio with its 95% C.I. and significance ( $p$ -level) computed using logistic regression with borreliosis, sex, and age as independent variables. The number of subjects reporting the results of a serological test for borreliosis and providing the information about their mental health disorders was 7,810. The association significant after the correction for multiple tests is printed in bold.

## 4 Discussion

The main finding of this study is the robust and specific effect of latent toxoplasmosis, or, more precisely, the presence of anamnestic titres of anti-*Toxoplasma* antibodies, on mental health symptoms and disorders. The most characteristic symptom associated with *Toxoplasma*-seropositivity was increased anxiety and the typical toxoplasmosis associated disorders were autism (OR=4.86), schizophrenia (OR=3.34), attention deficit hyperactivity disorder (OR=3.02), Asperger syndrome (OR=2.49), antisocial personality disorder (OR=1.81), obsessive compulsive disorder (OR=1.69), and general anxiety disorder (OR=1.57). On the other hand, the *Borrelia* seropositivity was associated only with symptoms of depression (n.s. after the correction for multiple tests) and with major depression (OR=1.65), despite that the size of the population tested for borreliosis was more than two times larger than the size of the population tested for toxoplasmosis. Moreover, the negative effects of the *Borrelia* seropositivity were observed in *Toxoplasma*-seropositive, but not in *Toxoplasma*-seronegative subjects.

The association of toxoplasmosis with schizophrenia has been confirmed by many studies, for reviews see (Sutterland et al., 2015; Torrey et al., 2012) and similar association has been documented for obsessive compulsive disorder (Flegr and Horáček, 2017; Miman et al., 2010; Xiao et al., 2010) and anxiety disorder (Gale et al., 2014). The association between toxoplasmosis and autism has been suggested on the basis of three case-controls studies and also on various indirect evidence, for review see (Flegr and Escudero, 2016; Prandota et al., 2015). The lack of association between toxoplasmosis and major (unipolar) depression and panic disorder is in agreement with most of published data (Flegr, 2015b; Flegr and Hodny, 2016; Gale et al., 2014; Sutterland et al., 2015). As far as we know, the strong association of toxoplasmosis with attention deficit hyperactivity disorder, and also the relatively strong association with antisocial personality disorder have not been reported. In contrast, we did not confirm the association of toxoplasmosis with bipolar disorder, which has been reported in about ten studies, for review and metaanalysis see (Sutterland et al., 2015).

The number of associations of mental health problems with *Borrelia* seropositivity was much lower than with *Toxoplasma* seropositivity, for review see (Fallon and Nields, 1994). This seems to agree with the situation existing in literature. Pubmed search for toxoplasm* AND (psychiatry* OR mental*) finds 563 hits while the search for borrel* AND (psychiatry* OR mental*) gives only 53 hits, mostly case reports or descriptions of neuropsychiatric effects of neuroborreliosis. It must be reminded that nearly a third of participants of our survey, which had been self-selected on the basis of their interests about sexual topics, i.e., probably randomly with respect to health problems or probability of being examined for vector-transmitted pathogens, were able to provide information about the results of their serological tests on borreliosis. The incidence of borreliosis is growing in the Czechia (and Europe), however, even after performing the extrapolation of the 1990-2010 trend it is still lower than 80 cases per 100 000 inhabitants (Zeman and Benes, 2013). This means that most of the seropositive participants of the study have not been diagnosed with acute or post-acute borreliosis or even neuroborreliosis. Our results therefore suggest that, possibly except a small sub-population of subjects who are also infected with *Toxoplasma*, people infected with *Borrelia* typically express no mental health problems. One large study showed increased seroprevalence of borreliosis in Czech psychiatric patients (33 %) compared to matched healthy control subjects (19 %) (Hajek et al., 2002) and one recent study performed in Poland demonstrated higher incidence of depressive disorders in 75 patients with neuroborreliosis (50.7%) and 46 patients with Lyme arthritis (39.1%) (Oczko-Grzesik et al., 2017). However, these groups of patients could represent small, atypical sub-populations of *Borrelia*-infected subjects with some special genetic or non-genetic vulnerability for psychiatric problems such as coinfection with the *Toxoplasma*.

Our results show that the effects of toxoplasmosis and borreliosis are not additive. *Borrelia* had a strong effect on mental health (depressiveness) in *Toxoplasma*-infected subjects and only marginal or no effects on the mental health of *Toxoplasma*-seronegative subjects. Currently, we have no explanation for this modulatory (sensitizing) effect of toxoplasmosis. The candidate explanation for this effect could be an interaction between pathophysiological cascades induced by both pathogens. The influence of *T. gondii* on the development of psychiatric disorders is mediated both by an immune reaction of the brain to *T. gondii* and by the biochemical activity of the parasite itself. Interferon-gamma secreted in response to toxoplasmosis maintains this infection in a latent form because it induces astrocytes enzymes responsible for tryptophan degradation via the kynurenine metabolic pathway (Hunt et al., 2017; Silva et al., 2002). It results in both a lack of tryptophan, an amino acid essential for *T. gondii* replication, and increased levels of the final products of kynurenine pathway. These metabolites exert both neurotoxic (quinolinate) and pro-psychotic (kynurenic acid) effects (Elsheikha et al., 2016) and can also influence the neurotransmitter balance (Krause and Muller, 2012). However, through the medium of two genes analogical to the human gene for tyrosine hydroxylase, *T. gondii* also directly enhances dopaminergic activity that is critical for the development of schizophrenia, autism and other mental disorders (Flegr and Horáček, 2017; Sutterland et al., 2015; Torrey et al., 2012).

*Borrelia burgdorferi* infection is also associated with increased levels of proinflammatory cytokines including interferon-gamma. In addition, *Borrelia burgdorferi* surface glycolipids and flagella antibodies appear to elicit anti-neuronal antibodies (Bransfield, 2012). The proinflammatory cytokines, anti-neuronal antibodies and *Borrelia* lipoproteins can disseminate from the periphery and activate the microglia in the brain by engagement of Toll-like receptors and exacerbate the neuroinflammatin (Bransfield, 2012). Increased serum levels of nitric oxide and nitrotyrosine (resulting from excessive levels of protein nitration and lipid peroxidation) detected in patients with borreliosis can also amplify the inflammatory processes and induce neuronal apoptosis (Parthasarathy and Philipp, 2017; Ramesh et al., 2015). Given the more pronounced negative effects of the *Borrelia* seropositivity in subjects co-infected by *Toxoplasma* in our sample, it is possible to speculate, that the neuroinflammation induced by borreliosis represents the additional hit for subjects with latent toxoplasmosis, which increases the risk of mental disorders. The fact that borreliosis has a weaker effect on psychiatric morbidity might be due to the fact that in contrast to toxoplasmosis, Lyme disease is not associated with increased dopaminergic activity (Blum et al., 2017).

We can only speculate about the increased risk of *Borrelia* seropositivity in *Toxoplasma*-infected subjects. It is known, however, that latent toxoplasmosis has specific immunomodulatory effects on immune system of infected humans and mice (Flegr and Stříž, 2011; Kaňková et al., 2010) which could influence the risk and the course of other infections such as borreliosis. An alternative explanation could be based on the hypothesis of the existence of a vector shared by both pathogens. Despite the fact that the tick infestation is not considered to be a usual source of human *Toxoplasma* infections, *Toxoplasma* DNA is detected in 12.6% of the most common European tick, *Ixodes ricinus,* which has approximately the same rate of positive tests as for *Borrelia* (12.7%) (Sroka et al., 2009). The same study showed that the existence of both pathogens occurred in 2.3% of all ticks and in 3.8 adult females. The transmission of *Toxoplasma* by three species of ticks (*Dermacentor variabilis*, *D. andersoni*, and *Amblyomma americanum*) has been also documented previously (Woke et al., 1953).

## 5 DStrength and limits of the present study

Our study probably involves the largest ever population of *Toxoplasma*-tested participants–the usual size of populations in similar studies is at least one order of magnitude smaller. Our study was exploratory and hypothesis-free in the sense that all main mental health disorders have been analyzed and all results, both positive and negative, have been reported. This approach remediates the well-known problems of the drawer and the cherry picking of artifacts – the problems of reporting only positive or “interesting” results of studies.

The most serious limitation of the present study is that the participants (about 0.5 % inhabitants of Czechia) have been self-selected and therefore they probably do not represent a typical Czech population. The study primarily concerned sexual behavior and sexual preferences and therefore mainly the subjects who were interested in these topics took part in it and finished the whole 110 minutes questionnaire. However, there is no reason to suppose that the association between the *Toxoplasma* or *Borrelia* infections and mental health would differ between the subjects who voluntarily participate in the sex-related study and in the general population.

Another limitation of the present study was that the subjects self-reported their mental health status, including the incidences of particular mental health disorders, as well as their infection status itself. This is a tradeoff for being able to study the interaction between acquired infections and mental health problems on a large enough sample. Our previous analyses of a sample of 3,827 subjects who had been tested in our laboratory for *T. gondii* seropositivity and later registered as our internet volunteers showed that the information concerning toxoplasmosis is mostly (99.5%) correct (Flegr, 2017). Still, it is highly probable that a certain fraction of raters misreport their psychiatric diagnoses and especially their self-diagnoses. The results of Monte Carlo modeling, however, showed that stochastic errors caused by misdiagnosing and misreporting health status can result only in false negative results of a study, i.e., the failure to detect existing associations, not false positive results of a study, i.e., detecting non-existing associations (Flegr and Horáček, 2017).

We did not control for the effect of the Rh phenotype in the present study. It has been shown recently, however, that Rh negative women express increased, while Rh-positive women express decreased depression, obsession, and several other facets of neuroticism measured with the N-70 inventory (Šebánková and Flegr, 2017).

Given the cross-sectional design of this study, we cannot address the problem of causality. It is neither probable that mental health disorder could increase the risk of acquiring the infection, nor of reporting the *Toxoplasma* infection but not the *Borrelia* infection. However, it is possible that some unknown third factor, such as immunodeficiency, could increase both the risk of mental health disorders and the *Toxoplasma* infections. In the light of already published data and our knowledge of the biology of the studied neuropathogens, however, the most parsimonious interpretation of the observed association is the positive effect of the infections on the rate of specific mental health disorders.

## 6 Concluding remarks

Our results obtained in the cross-sectional study performed on a cohort of more than 3 300 subjects tested for toxoplasmosis and more than 7 800 subjects tested for borreliosis suggests that the former pathogen has strong effects on the rate of several severe mental health disorders, including psychoses. Borreliosis showed only a weak association with major depression and with subjectively reported depressive and anxiety symptoms; these effects, however, seem to affect mainly (or exclusively?) *Toxoplasma*-co-infected subjects. It is important to mention that the course of acute toxoplasmosis in immunocompetent subjects is mild and is considered insignificant from the clinical point of view, while the opposite is often true for borreliosis. The results of current study suggest that despite seemingly asymptomatic course of latent toxoplasmosis, *Toxoplasma* could play the privileged role in the etiology of mental disorders.

## Acknowledgments

We would like to thank Lincoln Cline for her help with the final version of the paper. The work was supported the Czech Science Foundation [Grant No. P407/16/20958].

